# The effects of modified sialic acids on mucus and erythrocytes on influenza A virus HA and NA functions

**DOI:** 10.1101/800300

**Authors:** Karen N. Barnard, Brynn K. Alford-Lawrence, David W. Buchholz, Brian R. Wasik, Justin R. LaClair, Hai Yu, Rebekah Honce, Stefan Ruhl, Petar Pajic, Erin K. Daugherity, Xi Chen, Stacey L. Schultz-Cherry, Hector C. Aguilar, Ajit Varki, Colin R. Parrish

## Abstract

Sialic acids (Sia) are the primary receptors for influenza viruses, and are widely displayed on cell surfaces and in secreted mucus. Sia may be present in variant forms that include *O-*acetyl modifications at C4, C7, C8, and C9 positions, and *N*-acetyl or *N*-glycolyl at C5. They can also vary in their linkages, including α2-3 or α2-6-linkages. Here, we analyzed the distribution of modified Sia in cells and tissues of wild-type mice, or in mice lacking cytidine 5’-monophosphate-*N*-acetylneuraminic acid hydroxylase (CMAH) enzyme that synthesizes *N-*glycolyl modifications (Neu5Gc). We also examined the variation of Sia forms on erythrocytes and saliva from different animals. To determine the effect of Sia modifications on influenza A virus (IAV) infection, we tested for effects on hemagglutinin (HA) binding and neuraminidase (NA) cleavage. We confirmed that 9-*O*-acetyl, 7,9-*O*-acetyl, 4-*O-*acetyl, and Neu5Gc modifications are widely but variably expressed in mouse tissues, with the highest levels detected in the respiratory and gastrointestinal tracts. Secreted mucins in saliva and surface proteins of erythrocytes showed a great degree of variability in display of modified Sia between different species. IAV HA from different virus strains showed consistently reduced binding to both Neu5Gc and *O-*acetyl modified Sia; however, while IAV NA were inhibited by Neu5Gc and *O*-acetyl modifications, there was significant variability between NA types. The modifications of Sia in mucus may therefore have potent effects on the functions of IAV, and may affect both pathogens and the normal flora of different mucosal sites.

**IMPORTANCE:** Sialic acids (Sia) are involved in many different cellular functions and are receptors for many pathogens. Sia come in many chemically modified forms but we lack a clear understanding of how they alter the interactions with microbes. Here we examine the expression of modified Sia in mouse tissues, on secreted mucus in saliva, and on erythrocytes, including those from IAV host species and animals used in IAV research. These Sia forms varied considerably between different animals, and their inhibitory effects on IAV NA and HA activities and on bacterial sialidases (neuraminidases) suggest a host-variable protective role in secreted mucus.

## INTRODUCTION

Sialic acids (Sia) are a family of nine-carbon monosaccharides that often serve as terminal residues of carbohydrate chains. They are present at high levels on cell membrane glycoproteins and glycolipids, as well as on secreted glycoproteins and mucus at all mucosal surfaces (**Fig. 1A**) (1, 2). Sia are key mediators of many normal cell and tissue functions through a wide variety of highly regulated cell-cell interactions during both development and homeostatic processes, where they may be bound by cellular receptors and members of the selectin family (3, 4). Their ubiquitous presence on cells, tissues, and mucosal surfaces also make Sia a key point of contact for commensal microbes and for invading pathogens including viruses, bacteria, and parasites (3, 5, 6).

**Figure 1.**
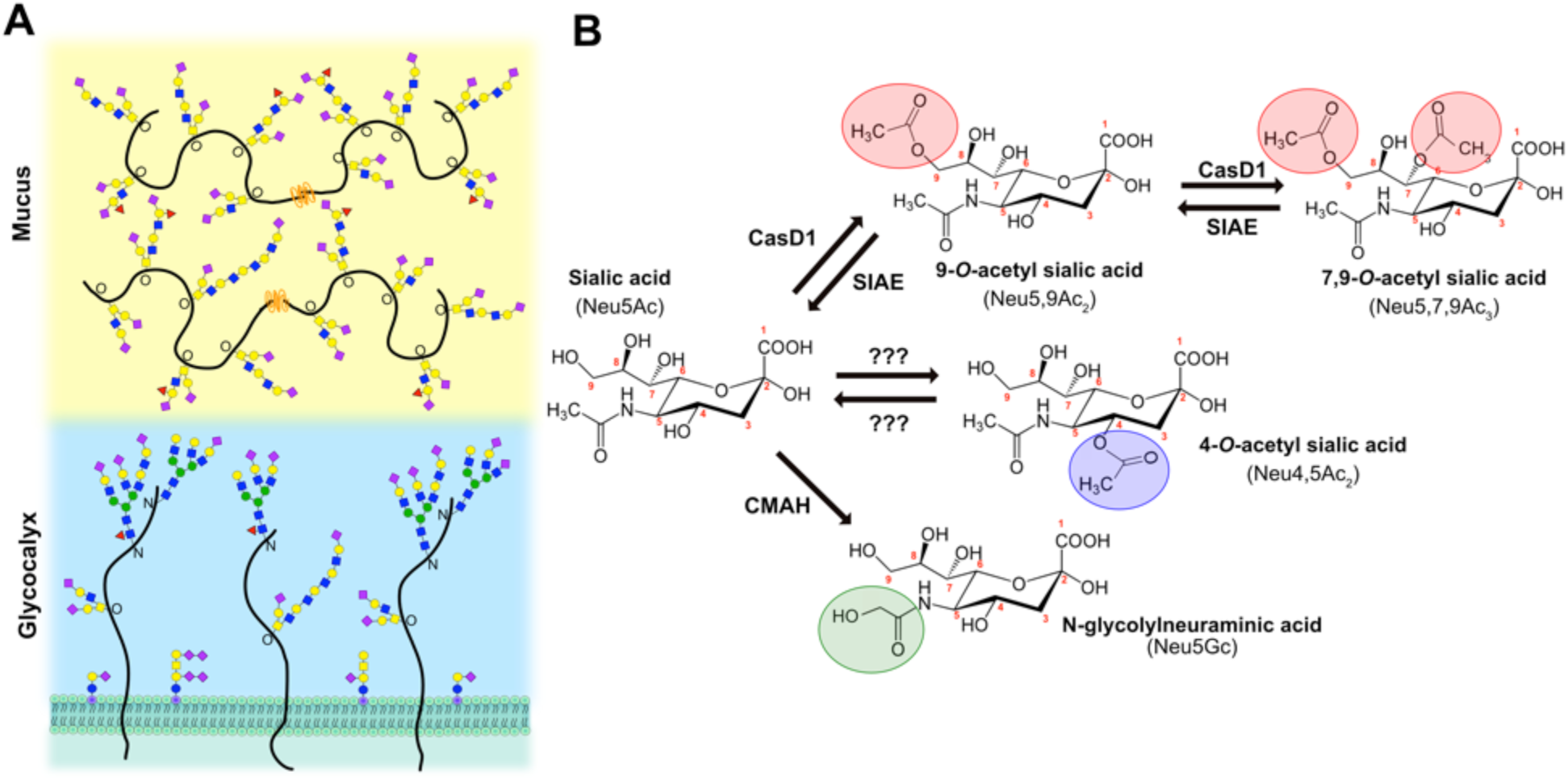
**A)** Sialic acids (purple diamonds) terminate glycan chains on glycolipids and glycoproteins as part of the glycocalyx on the surface of cells. They can also terminate glycans on secreted glycoproteins, like mucins, that are an important component of the protective mucosal barrier in gastrointestinal and respiratory tissue. **B)** Sialic acid (*N*-acetylneuraminic acid, Neu5Ac) can be modified by the addition of O-acetyl modifications at the C-4, 7, and 9 positions, or by the hydroxylation of the *N-*acetyl group at C-5 to form *N*-glycolylneuraminic acid (Neu5Gc) by the enzyme cytidine 5’-monophosphate-N-acetylneuraminic acid hydroxylase (CMAH). The sialate *O-*acetyltransferase, CasD1, adds acetyl groups at C-7 from which it migrates to the C-9 position (Neu5,9Ac_2_) under physiological conditions. This can allow for an additional acetyl group to be added by CasD1 to C-7 (Neu5,7,9Ac_3_). The sialate *O-*acetylesterase, SIAE, can remove these acetyl modifications, restoring the unmodified Neu5Ac form of sialic acid. *O-*acetyl modifications can also be added at the C-4 position by a specific 4-*O-*acetyltransferase (Neu4,5Ac_2_) and removed by a 4-*O-*acetylesterase. However, the genes for these enzymes have not yet been identified.

Sia are a highly diverse family of molecules that may be present as more than 50 structurally and chemically distinct modified variants. These are formed from the basic structure of the *N*-acetylneuraminic acid (Neu5Ac) by the addition of chemical groups at various positions on the pyranose ring or the glycerol side chain. Those modifications may include *N*-glycolyl and/or *O*-linked acetyl, sulfo, methyl, and lactyl groups, among others (1, 2, 7). Many different enzymes and pathways introduce these chemical modifications and some can be removed by regulatory enzymes. The different modified Sia are often themselves substrates for modifying enzymes and transferases, so that each modification may alter the synthesis of other modified forms. This therefore leads to complex patterns and mixtures of modified Sia forms, with significant variation in both the levels and specific combinations of modifications in different hosts, tissues, and under different physiological conditions (4, 8–11).

### Sialic acid modifications

Common chemical additions seen in vertebrates include ester-bonded *O*-acetyl (*O*-Ac) modifications to C-4, 7, 8, and/or 9 positions, resulting in a variety of combinations of Sia forms including Neu4,5Ac_2_, Neu5,9Ac_2_, Neu5,7,9Ac_3_ Sia, as well as their *N*-glycolylneuraminic acid (Neu5Gc) analogs with the same *O-*acetyl modifications (**Fig. 1B**). Neu5Gc is produced from Neu5Ac by the activity of cytidine 5’-monophosphate-N-acetylneuraminic acid hydroxylase (CMAH) in the cytoplasm of cells, and this enzyme is missing or inactive in some animals, including humans (9, 12). The addition of *O-*Ac to the C7 and/or C9 positions is mediated by the sialylate *O-*acetyltransferase enzyme, Cas1 domain containing 1 (CasD1), which has been suggested to add an *O-*acetyl group to the C7 position, from which it would migrate to the C8 and C9 position under physiological conditions, allowing the possibility of the addition of another *O-*acetyl group to C7 (7, 13, 14). The regulatory processes that control the number or positions of acetyl groups have not been well defined, although distinct differences in expression of 7,9-*O-*Ac and 9-*O-*Ac Sia have been reported in mouse and human tissues, chicken embryos, and in some other animals (7). CasD1 uses acetyl-CoA to modify Sia in the activated CMP-Sia form before it is added to the glycan chain, and likely has a preference for CMP-Neu5Ac as a substrate and is less active on CMP-Neu5Gc (14). The sialate *O-*acetylesterase (SIAE) enzyme can remove the 7,9-*O-* and 9-*O-*Ac modifications, although its activities and roles are not well understood (17–20). The 4-*O*-Ac Sia is produced in some tissues of many animals by a distinct sialate 4-*O*-acetyltransferase that is also likely expressed in the Golgi compartment; however, the gene for this enzyme has also not yet been identified (21–23).

The 7, 8, and/or 9-*O*-Ac Sia appear to be present at low levels - a few percent or less - in the cell-associated Sia on many cultured cells, but may be present at higher levels (10 to 50%) in Sia on the secreted mucus of various animals and on erythrocyte-associated glycans (15, 24). However, the expression, distribution, and regulations of these modified Sia are not well documented, nor do we understand their impact on pathogens, host homeostasis, and normal microbiota at different mucosal sites (9, 15, 16, 25).

### Modified sialic acid and pathogen interactions

Many pathogens interact with Sia on host cells at various stages in their infection cycles, including various viruses, bacteria, and parasites (3, 5). The densely expressed Sia within various mucus layers on mucosal surfaces also act to bind incoming pathogens and likely regulate both the release and transmission of pathogens (26, 27). Many pathogens therefore express proteins that attach to Sia, as well as expressing receptor-modifying enzymes such as sialidases (neuraminidases) that remove the Sia from the underlying glycan. Bacterial adhesins and toxins may recognize Sia on the surface of cells, and many bacteria can also use Sia as a metabolic carbon source after release through the activity of neuraminidases, and uptake into the cell by Sia transporters (3, 28–31). These bacterial interactions with Sia are potentially affected by chemical modifications (30, 32). Both enveloped and non-enveloped viruses may also bind Sia as primary receptors or co-receptors for cell recognition and infection, although only the enveloped viruses appear to express neuraminidases or sialate *O-*acetyl esterases, possibly to reduce aggregation of viral particles during budding (5, 33). For some viruses, Sia modifications are required for infection as viral proteins specifically bind to modified Sia – examples include human coronavirus OC43 and HKU1, and influenza C and D viruses, which all require 9-*O*-Ac Sia for cell infection (34, 35).

Significant effects of different Sia modifications on the binding of pathogens or the activities of their sialidases (neuraminidases) have been suggested, but in general these are still not well understood. Influenza A viruses (IAV) use Sia as primary receptors for host recognition and cell entry through the activity of two surface glycoproteins that interact with Sia, hemagglutinin (HA) and neuraminidase (NA). HA is a trimeric protein that binds Sia to initiate the endocytic uptake of the virus by the cell, leading to fusion between the viral envelope and the endosomal membrane after exposure of the virus to low pH (36). NA is a sialidase which cleaves Sia from the mucus, cell surface, and from viral glycoproteins, allowing the virus to penetrate through mucus to the epithelial cells and reducing the aggregation of virions after budding from the surface of cells (37). Previous studies have suggested that 7,9-*O*- and 9-*O-*Ac Sia are expressed on cells or in tissues of many IAV host species and there is some evidence that these modifications may be inhibitory for NA activity and HA binding (38, 39). However, the specific effects of 7,9-*O-* and 9-*O-*Ac on the binding of Sia by HA or the cleavage of Sia by IAV NA have not been examined in detail, and it is unclear whether these changes influence infection efficiency or viral shedding. For example, modification of HA binding might influence the attachment of virus to cells or to mucus, while inhibition of NA cleavage of *O-*acetyl modified Sia may lead to virus being trapped in the mucus and cleared, reducing the efficiency of infection.

The difference between Neu5Gc and Neu5Ac has been found to influence the tropism of several different viruses, as well as some bacterial toxins (3, 5). Indeed, it has been proposed that the loss of the *CMAH* gene in humans was an adaptive response to pathogen pressures (40). Neu5Gc is highly expressed in some tissues of IAV natural host species, including pigs and horses, and is also present in the tissues of mice and guinea pigs, which are frequently used as animals models (9, 12, 41). Neu5Gc has been seen to prevent binding of the HAs of some IAV, particularly in human-adapted strains (39, 42), but the effects on NA have not been well characterized. Nevertheless, examination of swine IAV isolates found distinct strain differences in their ability to cleave Neu5Gc by sialidase activity which were generally lower than against Neu5Ac (43).

Here we define the expression of 7,9-*O-*Ac, 9-*O-*Ac, and Neu5Gc modified Sia in the mucus, saliva, and on erythrocytes of different IAV host animals, as well as on the tissues and secreted mucus of mice. We also test the effects of these modifications on HA binding and NA activity of different strains of IAV, as well as their potential to alter virus infection.

## RESULTS

### Distribution of modified Sia in mouse tissues

Previous research on display of modified Sia in animal tissues and cultured cells have shown varying distributions of 7,9-*O-*Ac, 9-*O-*Ac, and 4-*O*-Ac Sia depending on the animal and the tissue examined (15, 16). Mice are an important model species for biomedical research, and some tissues have previously been screened for *O-*Ac display using probes derived from viral glycoproteins (virolectins) and by other methods (15, 16). We examined the distribution of modified Sia in a variety of tissues of wild-type (WT) C57/BL6 mice. Both 9-*O*- and 7,9-*O-*Ac were found throughout the lung and trachea as well as in the tracheal sub-mucosal glands that produce most of the mucus (**Fig. 2A**). These modified Sia were also found throughout the GI tract, with staining associated with epithelial cells, goblet cells, and associated mucus layers of the gastrointestinal tissues, including the stomach, small intestine, and colon (**Fig. 2B**). Interestingly, 9-*O-*Ac appeared to be present in higher amounts in most tissues, including salivary gland and esophagus, while the 7,9-*O-*Ac staining was minimal. However, 7,9-*O-*Ac did stain stronger than 9-*O-*Ac in stomach-associated mucus, and on tracheal epithelial cells. This seems to indicate that while the same enzymes (CasD1 and SIAE) are considered to control the presence of these modifications, the expression of 9-*O-* and 7,9-*O-*Ac are differentially regulated in individual tissues. The 4-*O*-Ac Sia showed high levels of probe binding in the colon, primarily in the mucus, and there was also some expression on the mucosal surfaces in the stomach, small intestine (jejunum) and trachea, as well as on cells within the red pulp of the spleen (**Fig. 2A, C; Supplementary Fig. 1**).

**Figure 2.**
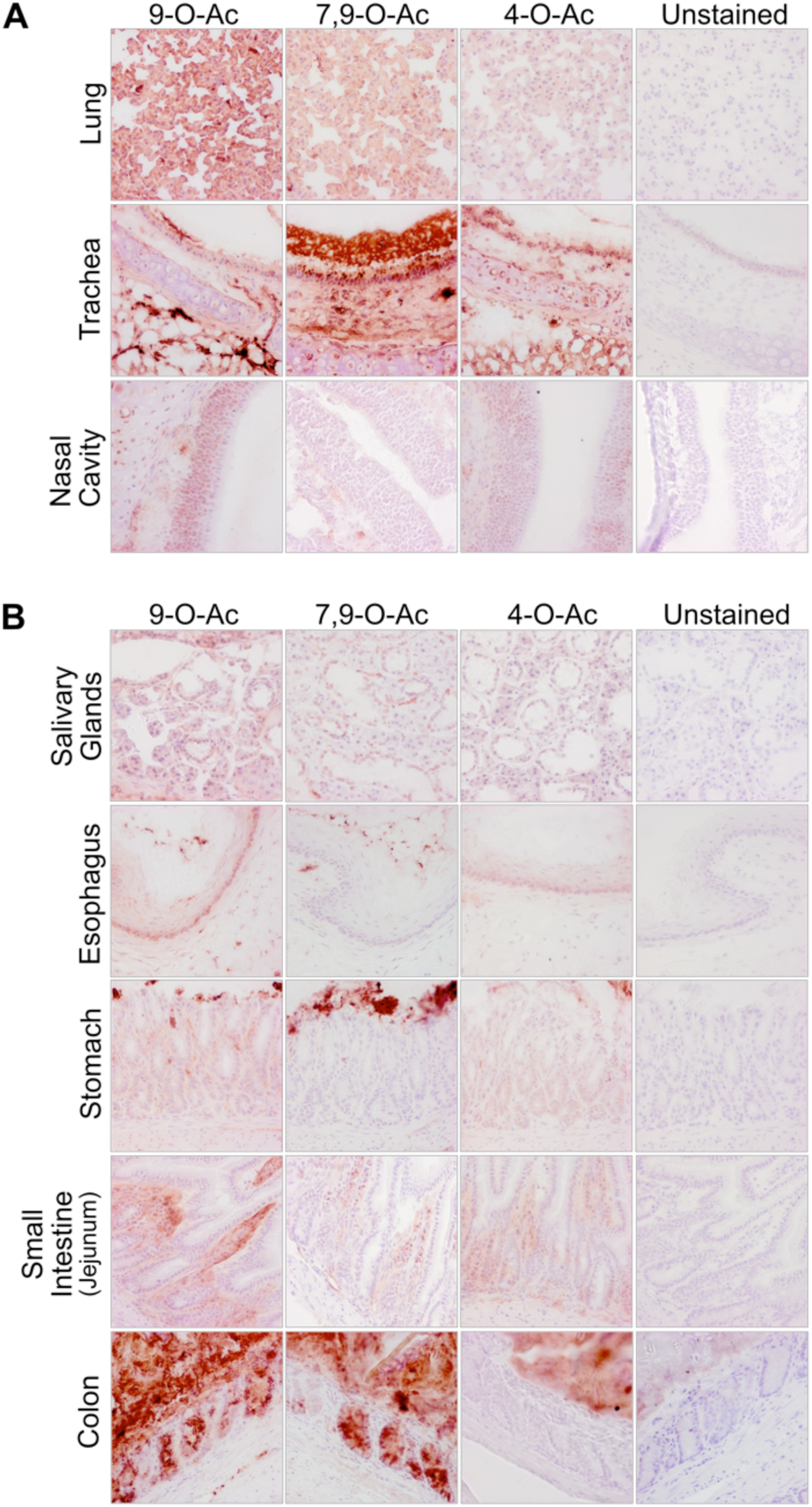
Expression of *O*-acetylated Sia varies between tissues in wild-type C57BL/6 mice Frozen tissue sections from respiratory tissues (**A**) and gastrointestinal tissues (**B**) were stained using virolectins derived from the hemagglutinin esterases (HE-Fc) of various nidoviruses with high specificity for the different *O-*acetyl modified Sia forms. Sections were counterstained with hematoxylin and imaged at 40x magnification. Stains from further tissues are shown in **Supplemental Figure 1**.

The virolectin probes used are sensitive, but did not reveal the quantity of each of the modified Sia forms present. To determine the relative amounts of different Sia forms, we tested tissues from mice using 1,2-diamino-4,5-methylenedioxybenzene (DMB) labeling of the Sia and analysis with high performance liquid chromatography (HPLC), under conditions that reveal the amounts of Neu5Ac and Neu5Gc, and preserve most of the *O-*acetylation of the Sia (45). The WT mice showed varying levels of Neu5Gc (**Fig. 3A; Supplemental Table 1**), while, as expected, the CMAH^-/-^ mice showed only Neu5Ac in all tissues, similar to amounts reported previously for some of those tissues (44) (**Supplemental Table 2**). The lack of Neu5Gc in CMAH^-/-^ mice was also seen in GI tissues, indicating that Neu5Gc from dietary sources was not detectably being taken up by these mice, as is seen in humans who eat a diet containing that Neu5Gc (46). In the WT mice, tissues showed a great deal of variability in Neu5Gc expression. Most tissues had around 50–60% Neu5Gc; however, some tissues, including the liver, had higher levels of over 70%, while the brain and salivary glands had only 10%. All *O-*acetyl Sia variants combined comprised between 2 and 16% of the Sia in most tissues, with the majority being 9-*O-*Ac (1–9% of total Sia) (**Fig. 3B; Supplemental Table 1**). There was about 1.5 to 3 fold higher levels of *O-*acetyl Sia in most tissues of the CMAH knock-out mice (**Fig. 3C; Supplemental Table 2**), as has been reported previously (44). The levels of 4-*O*-Ac Sia were generally low, making up ∼2% of the Sia in small intestine (duodenum) and ∼1% in spleen, testes, and esophagus. Mouse colon samples showed the highest levels of total *O-*acetylation, with ∼17% of Sia having one or more *O-*acetyl modification, again primarily 9-*O-*Ac. Given the patterns seen using the HE-Fc virolectin staining, the 7,9-*O-* and 9-*O-*Ac forms must be present at high levels within or on certain cell sub-populations, as well as within mucus or mucus-secreting cells. For example, the high levels of 7,9-*O-* and 9-*O-*Ac found in the mouse colon were most likely associated with secreted mucus as most of those modified Sia were present in goblet cells (**Fig. 2B**). However, the differences seen between probe binding and the ratios of the different modified Sia within the total Sia underscore the importance of quantifying the different forms.

**Figure 3.**
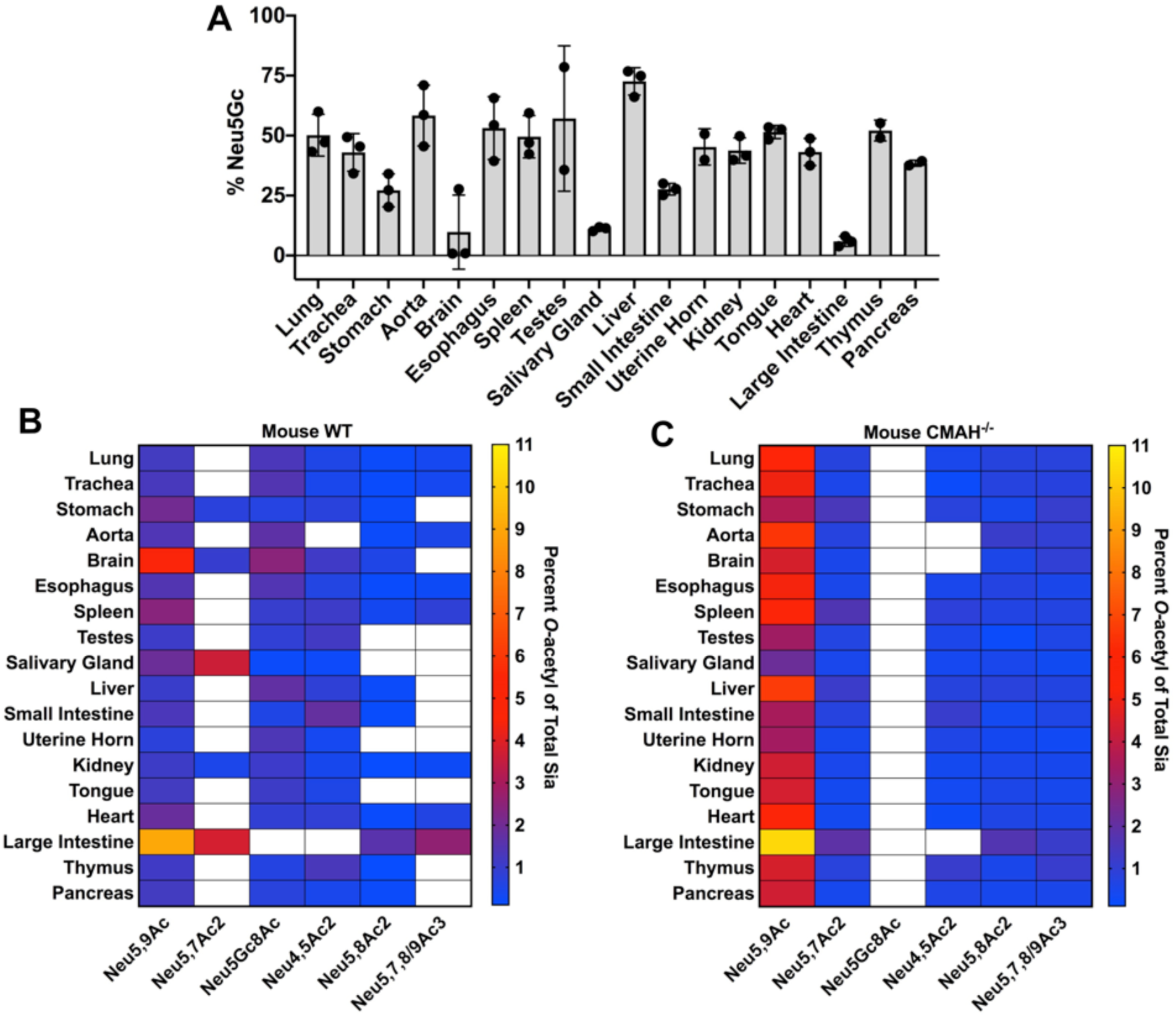
Sia modifications vary by tissue in wild-type C57BL/6 mice, and loss of Neu5Gc in CMAH^-/-^ mice leads to an increase in *O-*acetylation across tissues. Total Sia was measured from tissue samples using HPLC analysis to determine relative Sia quantities. (**A**) Neu5Gc levels were measured across tissues, showing highly specific expression. *O-*Acetylation in WT (**B**) and CMAH ^-/-^ (**C**) mice is given as a heat map showing variation across tissues. White squares indicate when a Sia form is below detection. Values are given as the percentage of total Sia collected from tissue samples. Sample size for each tissue was three individual mice (n=3) of each mouse strain with average values for total sialic acid content given in **Supplemental Tables 1 and 2**.

### Analysis of modified Sia in saliva, mucus, and on erythrocytes

It has been previously reported that human colonic mucin is highly enriched in 9-*O-*Ac Sia, which may regulate the activity of some sialidases and Sia transporters of the gut microflora (11, 32, 47, 48). Strong staining for 9-*O-*Ac in human respiratory tissues, and also within the submucosal glands of human respiratory tissue have also been reported, indicating that mucus from these glands could be enriched in *O-*acetylated Sia (47, 48). To determine if human respiratory mucus was enriched in 7,9-*O-* and 9-*O-*Ac, secreted mucus from primary normal human bronchial epithelial cells (NHBE) as well as conditioned media from human alveolar basal epithelial adenocarcinoma A549 cells, were analyzed by HPLC to determine Sia composition. We found that the secreted proteins in mucus from NHBE cells and A549 cells conditioned media contained primarily unmodified Neu5Ac with ∼1–2% of 9-*O-*Ac and no detectable levels of 7,9- *O-*Ac (**Table 1**). This indicates that secreted mucus from these respiratory cells in culture are not enriched for *O-*acetyl modifications, which differs from previous reports for colonic mucin (11, 47).

**Table 1.** A549 conditioned media and collected mucus from NHBE cultures were analyzed for total sialic acid content using HPLC analysis. Table shows the proportion of total Sia for each variant given as a percentage, with the sum of all *O-*acetyl forms combined given in the far right column. If a Sia form was below detection, this is indicated by b/d. Percentages are an average of multiple conditioned media collections from A549 cells (n=4) and multiple NHBE donors (n=4).

| Cells | Source | Neu5Gc | Neu5Ac | Neu5,9Ac | Neu5,7Ac<sub>2</sub> | Neu5,8Ac<sub>2</sub> | Neu5,7,8/9Ac<sub>3</sub> |
| --- | --- | --- | --- | --- | --- | --- | --- |
| A549 | Conditioned media* | b/d | 98.16 | 1.84 | b/d | b/d | b/d |
| NHBE | mucus | b/d | 98.6 | 1.40 | b/d | b/d | b/d |

\*: collected from A549 bronchial epithelial cells conditioned media, contains mucins

To look more broadly at the possible range of modified Sia present in secreted mucus in different animals, we examined saliva from a number of influenza host species, including human, pig, horse, and dog (**Fig. 4A,B**). While the proteins in saliva differ from those found in respiratory mucus, they do contain some of the same heavily glycosylated proteins including mucins like MUC5B (49–51). Human saliva was similar to the secreted mucus from NHBE and A549 cells in containing primarily Neu5Ac with little 9-*O-*Ac Sia, and the composition of dog saliva showed a similar profile. However, most other animals showed far more diversity in their Sia profiles, with both mice and horses having enrichment for several different *O-*acetyl modifications. Laboratory mouse saliva contained a combined ∼17% *O-*acetylated Sia in the forms of 7-*O-, 8-O-*, and 9-*O-*Ac, while horse saliva contained ∼10% 4-*O-*Ac as well as ∼19% of other *O-*acetyl Sia variants combined. Pig saliva were unique among the IAV hosts examined in having ∼90% of total Sia being of the Neu5Gc form. The diversity of modifications seen in mouse, horse, and pig saliva may have a strong influence on any Sia-binding pathogens, including influenza viruses, as well as on commensal bacterial communities in different species.

**Figure 4.**
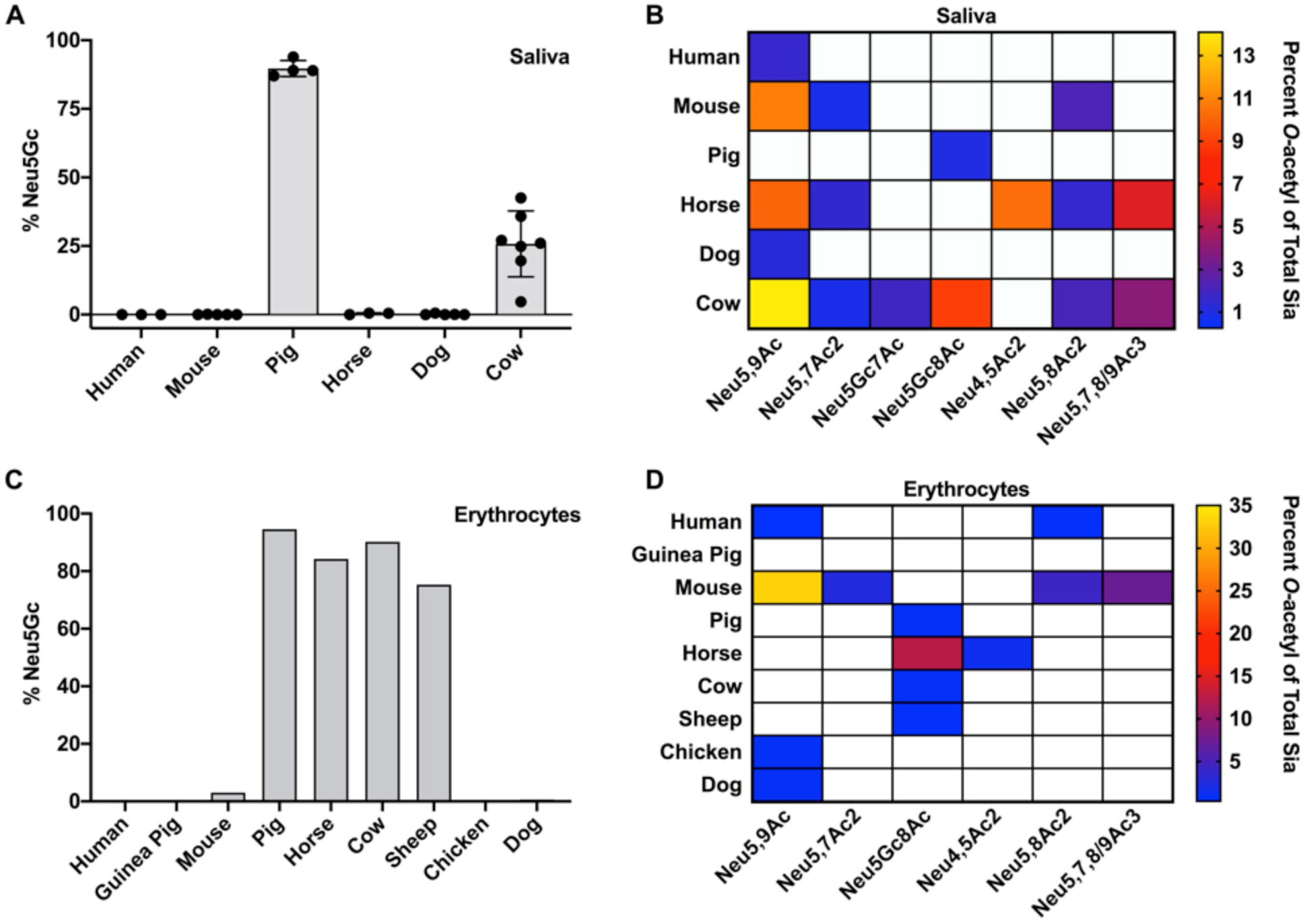
Total Sia of saliva (**A,B**) and erythrocytes (**C,D**) was collected via acid hydrolysis and analyzed using HPLC. *O-*Acetyl Sia percentages are given as a heat map with white squares indicating when a Sia form is below detection. Values are given as the percentage of total Sia collected from tissue samples. Saliva sample sizes (n = number of individuals of each species): human (n=3), mouse (n=5), pig (n=4), horse (n=3), dog (n=5), cow (n=7). Erythrocyte sample sizes (n = number of individuals of each species): human (n=2), guinea pig (n=1), mouse (n=2), pig (n=1), horse (n=1), cow (n=1), sheep (n=1), chicken (n=1), dog (n=2).

Erythrocytes (red blood cells, RBCs) express high levels of sialylated surface molecules, primarily on glycophorins, and are used in IAV research to study the interactions of HA binding specificity, determining viral titer through the hemagglutination assay, and inhibition of hemagglutination by antibodies (HAI assay) (52, 53). It has long been known that IAV varies in hemagglutination of RBCs from different species, at least in part due to differences in the Sia linkages present. The structures of HAs with Sia bound often suggest that modification of the C4, 5, 7, and/or 9 positions would influence IAV interactions with Sia. We found that chicken and guinea pig RBCs, which are often used to titer IAV virus and as the standard substrate for HAI assays, contained almost exclusively unmodified Neu5Ac, as did those from humans and dogs (**Fig. 4C,D**). In contrast, pig, horse, cow, and sheep RBCs contain high proportions of Neu5Gc, along with varying amounts of *O-*acetyl modifications. The high levels of Neu5Gc Sia present on the RBCs of these species had been previously reported, although not directly quantified as presented here (24, 54). The mouse erythrocytes tested here (from C57BL/6 mice) had the greatest diversity of modifications, with little Neu5Gc, around 50% Neu5Ac, and ∼50% of the Sia modified by 7, 8 and/or 9-*O*-acetylation, as previously reported (55).

### Effects of modified Sia on NA cleavage

Neuraminidases (sialidases) expressed by bacteria and viruses cleave Sia from oligosaccharides and glycoconjugates, and some have been shown to be affected by various Sia modifications (38, 42, 54, 56, 57). But little is known about the effects of modified Sia on IAV NAs from different strains. We examined the effects of Sia modifications on cleavage by several different IAV NAs, as well as on the activity of *Arthrobacter ureafaciens* neuraminidase (NeuA). Substrates used had high levels of 7,9-*O-* and 9-*O-*Ac Neu5Ac (bovine sub-maxillary mucin, BSM), Neu5Gc (horse RBCs), or unmodified Neu5Ac (chicken RBCs). IAV NA from a variety of different IAV strains (N1, N2, N3, N7, and N9) were expressed alone in cells (**Fig. 5A**) and recovered as purified VLPs (**Fig. 5B**) (58). These NA VLPs were first tested for enzymatic activity using a standard NA cleavage assay using methylumbelliferyl *N-*acetylneuraminide (MuNANA) as the substrate (**Fig. 5C**) (42). The NA-expressing VLPs were incubated with BSM or with RBCs, and the released Sia were collected and analyzed by HPLC. For BSM, HPLC profiles of total Sia were created to compare NA cleavage preferences (**Fig. 6A**). These profiles showed the Sia forms that were susceptible to NA cleavage and release while the non-released forms were considered to be resistant to NA. These HPLC profiles were then compared to the total Sia released chemically by acid hydrolysis, an unbiased method that removes all Sia forms present in the original sample (45). All of the viral NA and the bacterial NeuA showed the highest level of cleavage for unmodified Neu5Ac compared to any of the modified forms, as more Neu5Ac was present in the released profiles compared to chemical release. There was substantial variation in the cleavage activity against the modified Sia by the different viral NAs. N1 and N7 showed the lowest activities against any modified Sia, N3 and N9 had intermediate activities, while N2 and NeuA were active against the greatest number of modified forms, and most closely matching the chemical release profile. There was lower activity for mono-*O-*acetylated Sia (7-*O-, 8-O-*, and 9-*O-*Ac) by N1, N3, N7, and N9, while all NAs tested had lower activity against the di-*O-*acetylated Sia (7,8/9-*O-*Ac_2_) and mono-*O-*acetylated Neu5Gc forms. All the viral and bacterial NAs apart from N2 had several-fold lower activities on Neu5Gc alone, as seen in the smaller proportion of that Sia form released compared to the chemical release profile.

**Figure 5.**
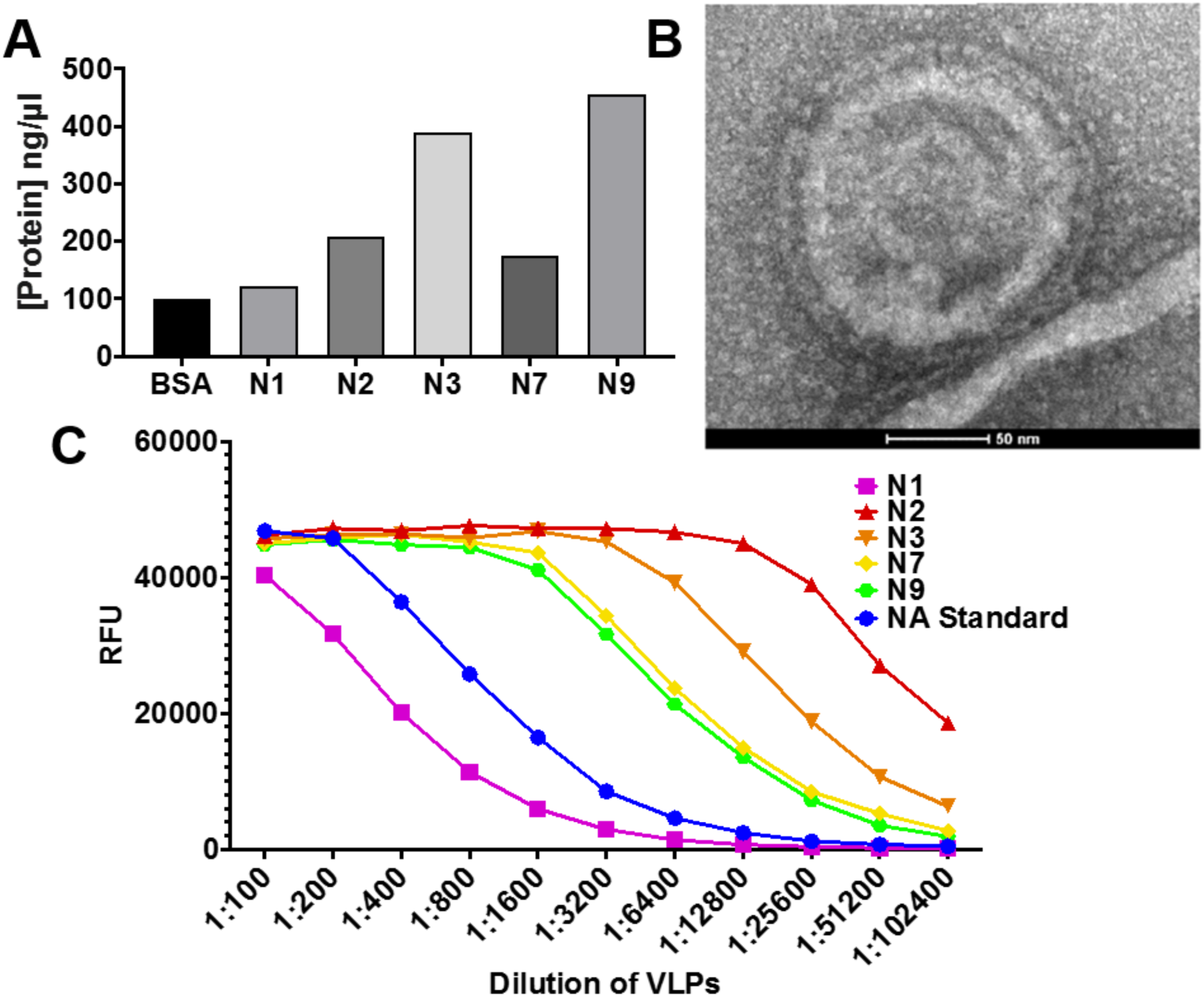
NA budding from mammalian cells. **A**) NA levels after control background subtraction from Coomassie Blue stain of NA protein expression in VLPs. **B**) TEM micrograph of a VLP expressing N2. **C**) Comparison of NA enzymatic activity using a MuNANA assay between different NA serotypes. One representative experiment is shown (n = 3).

**Figure 6.**
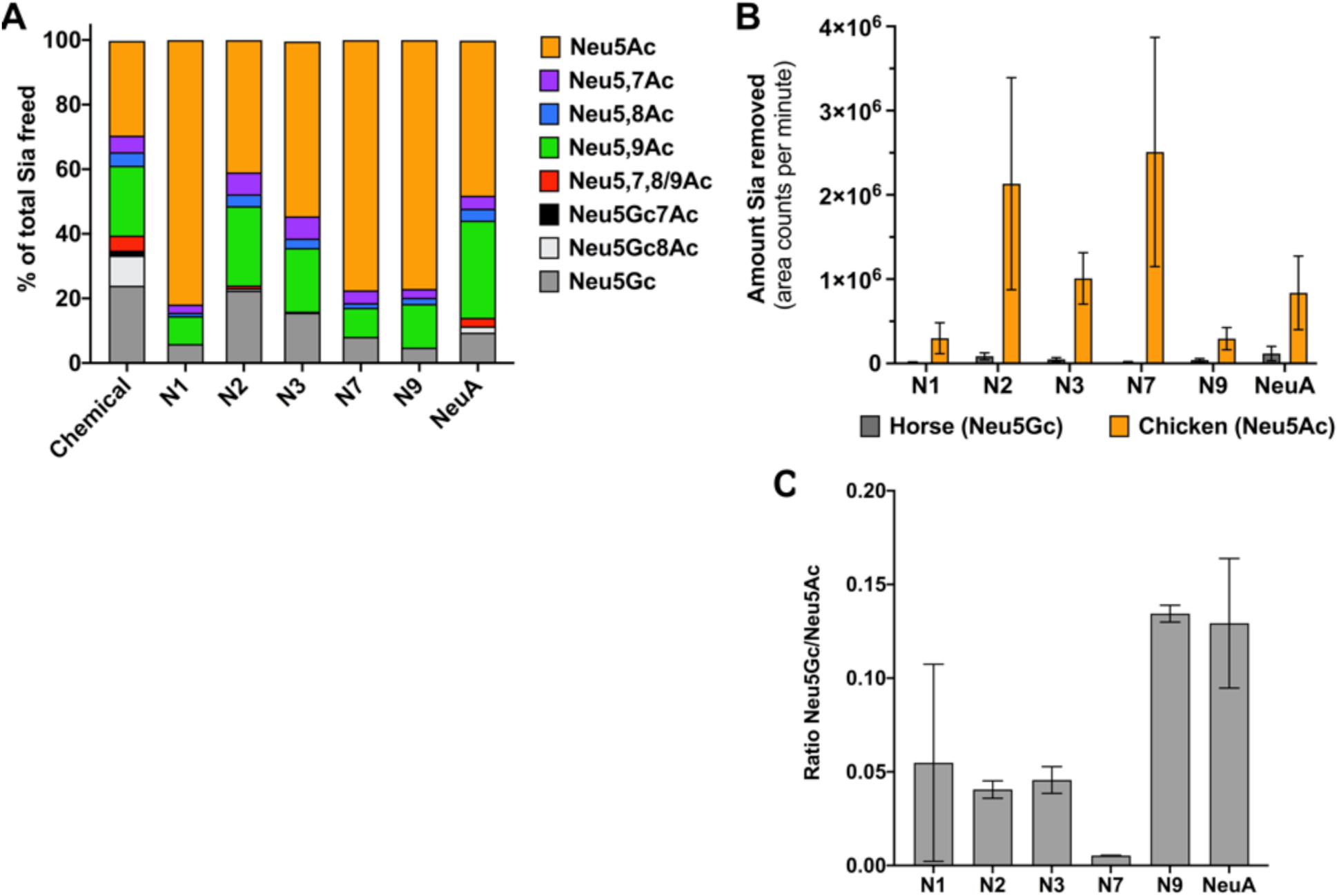
NA VLPs preferentially cleave unmodified Neu5Ac Sia and NA activity is inhibited by *O-*acetyl and Neu5Gc modifications. **A**) Bovine sub-maxillary mucin was treated with 1:100 NA VLPs or *Arthrobacter ureafaciens* NA (NeuA) for 4 hrs at 37°C and freed Sia was collected and analyzed using HPLC. The profiles of freed Sia were then compared to the profile of Sia removed chemically, a more unbiased approach. Profiles shown are the average of two independent experiments. **B,C**) Chicken erythrocytes (Neu5Ac) or horse erythrocytes (Neu5Gc) were treated with 1:100 NA VLPs for 4 hrs at 37°C and freed Sia was collected and total Sia removed was determined using HPLC. Average area counts per minute (area under curve of chromatogram) were used a measure of Sia removal to compare between Sia released form chicken and horse erythrocytes. Data shown is average of two independent experiments.

To further test the ability of these NA VLPs to cleave Neu5Gc compared to Neu5Ac, the amounts of Sia released from either horse RBCs (84% Neu5Gc) or chicken RBCs (99% Neu5Ac) were compared. All NAs showed significantly lower levels of Neu5Gc Sia released from horse RBCs compared to the amounts of Neu5Ac released from chicken RBCs (**Fig. 6B**). When compared directly, NA VLPs removed 5–12% of Neu5Gc compared to their activities on Neu5Ac (**Fig. 6C**). Again, variability was seen between NA VLPs here, with N7 having the lowest activity against Neu5Gc compared to Neu5Ac and N9 having the most. There was also variability in cleavage activity between NA from different strains as well, as seen in the variable amount of Neu5Ac removed by the NA VLPs in **Figure 6B**, but it is unclear whether this difference is due to the intrinsic activates of the NAs when expressed as VLPs or to innate differences in the specific activities of each NA enzyme, or both. It is clear, however, that *O*-acetyl and Neu5Gc modifications inhibit the activities of many different IAV NA and of NeuA, and that multiple modifications, such as di-*O-*acetyl modifications, or Neu5Gc that is also *O-*acetylated, were even more inhibitory, showing high resistance to most of the viral and bacterial NAs tested here.

### Effects of modified Sia on HA binding

The initiation of IAV infection requires HA glycoprotein binding to Sia to allow the virus to be taken up into the cell, and there appears to be a direct relationship between Sia binding affinity and infection (59). To clearly determine the effect of Sia modifications on HA binding, we examined the binding of soluble HA fused to a human IgG1 Fc (HA-Fc) to synthetic, biotinylated α2–6 linked sialosides of Neu5Ac, Neu5Ac9NAc, and Neu5Gc (60). Neu5Ac9NAc was used instead of 9-*O-*acetyl Neu5Ac (Neu5,9Ac_2_) due to the increased stability of the 9-*N*-Ac group (61, 62). Briefly, ELISA-grade 96 well plates were coated with HA-Fc derived from California/04/2009 H1N1 and Aichi/2/1968 H3N2 strains, then incubated with the synthetic biotinylated sialosides. Binding of the biotinylated sialosides to the HA-Fc was detected using a streptavidin-linked HRP probe as previously described (63, 64). Compared to Neu5Ac sialosides, both California/04/2009 H1 and Aichi/1968 H3 HA-Fcs had decreased binding to Neu5Gc and Neu5Ac9NAc (**Fig. 7A**). The addition of the *N-*acetyl group at C9 blocked most binding, while Neu5Gc showed only a low level of binding. We saw the same binding dynamics for other H1 and H3 HA-Fc, but the SNA lectin, which recognizes α2–6-linked Sia, bound equally well to Neu5Ac and Neu5Gc, but not to Neu5Ac9NAc (**Fig. 7B**). This shows that the presence of *O-* (and in this case *N*-) acetyl modifications can inhibit many Sia-binding proteins.

**Figure 7.**
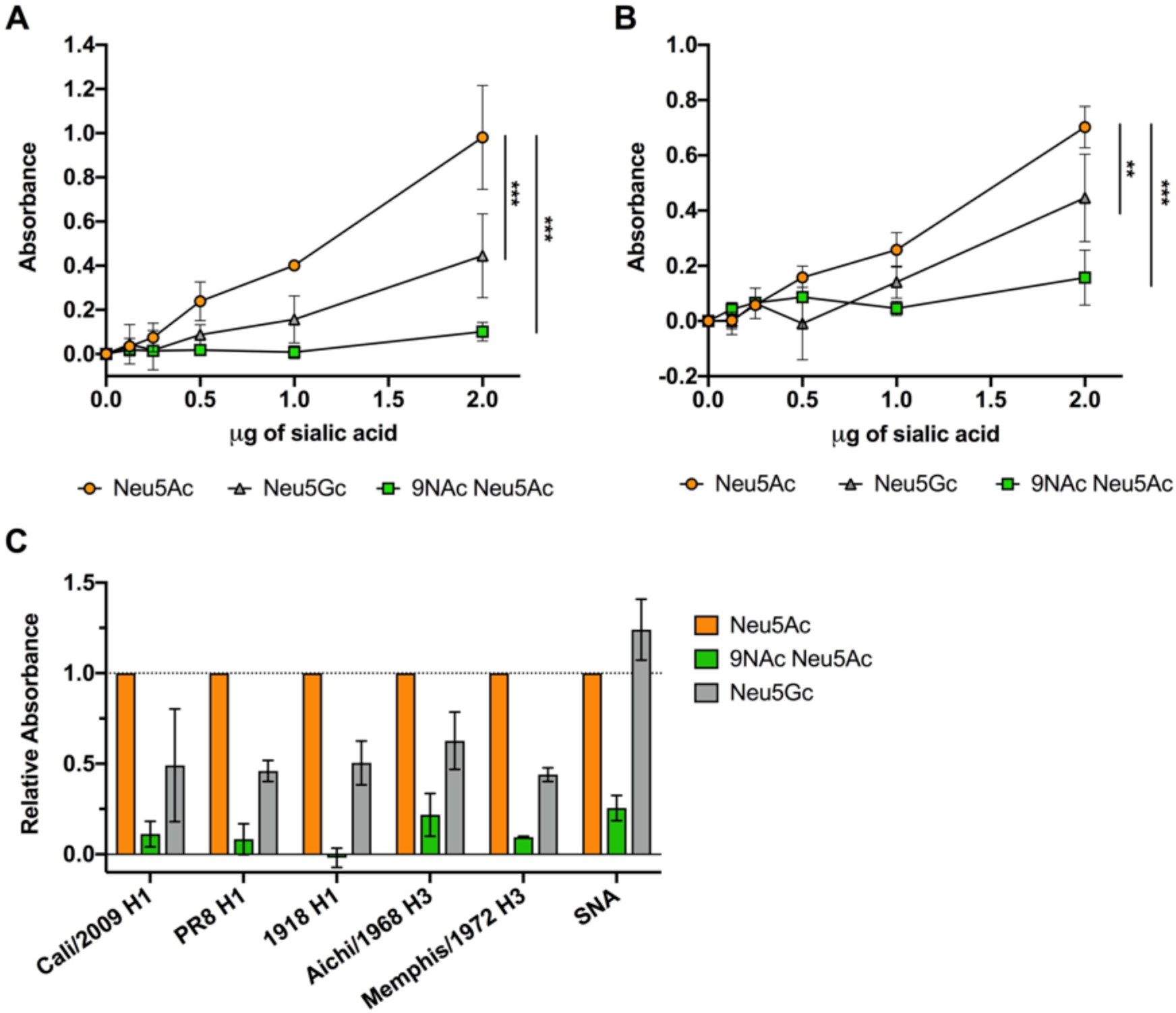
Soluble HA-Fc binding to synthetic sialosides showed decreased binding to modified Sia in an ELLA assay. **A,B**) Soluble HA constructs were developed by expressing HA proteins from different IAV strains fused to a human IgG1 Fc (HA-Fc). HA-Fc binding to synthetic α2–6-linked sialosides was assessed using an ELLA assay. Titration curves of sialoside binding by HA-Fc for (**A**) A/California/04/2009 H1N1 and (**B**) A/Aichi/2/1968 H3N2 were measured via colorimetric measurement. **C**) Sialoside binding for different H1 and H3 HA-Fc were determined using 2 µg of sialic acid. Lectin from *Sambucus nigra* (SNA), which specifically binds α2–6-linked Sia, was also included as a control. Data is shown as relative to HA-Fc binding to unmodified Neu5Ac. Data analyzed by 2-way Anova using PRISM software. Data is average of three independent experiments. **\*** = p-value ≤0.05; ** = p-value ≤0.01; *** = p-value ≤0.001.

### Effects of modified Sia on influenza A infection

While the low surface expression of 9-*O*- and 7,9-*O*-Ac on cells does not reduce IAV infection, viruses will also encounter modified Sia in mucus, which in many hosts and tissues has larger amount of these modifications. To determine how these Sia modifications can affect IAV infection, untreated BSM or BSM treated with esterase to remove 9-O-Ac and 7,9-O-Ac, were incubated with A/California/04/2009 (pH1N1), A/Puerto Rico/8/1934 (PR8 H1N1), and A/Victoria/361/2011 (Victoria H3N2) prior to infection of cells (**Fig 8A**). Both untreated BSM and esterase-treated BSM were inhibitory towards all three IAV strains, a trend towards higher inhibition by esterase-treated BSM, and significantly more inhibition for the PR8 H1N1 strain. This suggests that removing the 7,9-*O-* and 9-*O-*Ac from the Sia may have increased the virus binding to the mucin and inhibition of viral infection.

**Figure 8.**
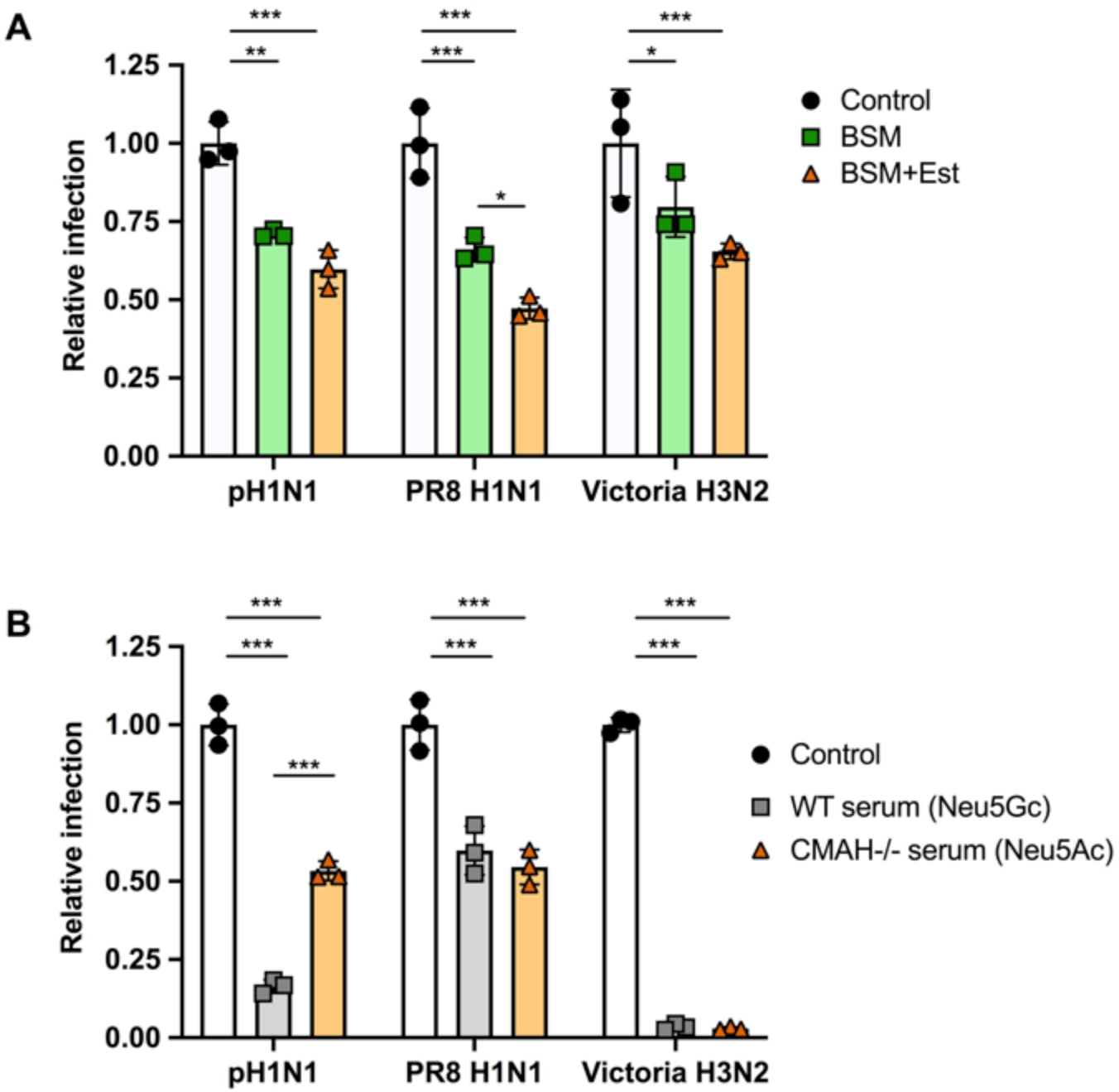
Virus infection is inhibited by mucin with trend for greater inhibition when *O-*acetyl groups are removed. Virus infection is also inhibited by serum, with no clear difference between Neu5Ac and Neu5Gc presence. **A)** A/California/04/2009 (pH1N1), A/Puerto Rico/8/1934 (PR8, H1N1), and A/Victoria/361/2011 (Victoria, H3N2) were mixed with 20 µg of BSM or BSM pre-treated with esterase active bovine coronavirus (BCoV HE-Fc) to remove *O-*acetyl modifications. This mixture was then used to infect cells at an MOI of 0.5 for 10 hrs. Infectivity was determined by flow cytometry analysis for NP positive cells. **B)** A/California/04/2009 (pH1N1), A/Puerto Rico/8/1934 (PR8, H1N1), and A/Victoria/361/2011 (Victoria, H3N2) were mixed with serum from either WT mice (Neu5Gc) or CMAH^-/-^ mice (Neu5Ac). This mixture was then used to infect cells at an MOI of 0.5 for 10 hrs. Infectivity was determined by flow cytometry analysis for NP positive cells. Data analyzed by 2-way Anova using PRISM software. **\*** = p-value ≤0.05; ** = p-value ≤0.01; *** = p-value ≤0.001.

Sia in sera have long been known to bind to influenza viruses, so that sera are often treated with neuraminidase as a “receptor destroying enzyme” prior to use in serological tests (65, 66). To specifically compare the effects of added Neu5Gc or Neu5Ac on the efficiency of infection, the same three IAV strains were incubated with mouse serum from either wild-type mice (>80% Neu5Gc) or CMAH^-/-^ mice (100% Neu5Ac) prior to inoculation of cells (**Fig. 8B**). In this case no specific trend was detected for the three viruses tested. Victoria H3N2 showed almost complete inhibition of infection by both sera, while PR8 H1N1 had a lower level of inhibition. Only pH1N1 showed a significant difference in response to the serums, with the WT mouse serum having a much stronger inhibitory effect compared to the CMAH^-/-^ serum. This suggests that Neu5Gc vs. Neu5Ac inhibition may vary by virus strain, and that likely depends on some combination of HA binding specificity and NA activity.

## DISCUSSION

Modified Sia are widely expressed within tissues and on mucosal surfaces of many animals, but with significant variation in the amounts of each modified Sia present (15, 16). Modified Sia are present at high levels on mucosal surfaces, including in GI and respiratory tissues, indicating that they are likely involved in tissue- and host-specific interactions with both pathogens and normal flora. While there have been suggestions that 9-*O*-Ac, 7,9-*O*-Ac, and Neu5Gc Sia might influence IAV infection by interfering with HA binding or NA activities, there is not a lot of direct evidence for their effects. Here we re-examine and define the tissue-specific and mucus expression of the 9-*O*- and 7,9-*O*-Ac-Sia as well as Neu5Gc, and to directly test their effects on IAV HA binding, NA activity, and infection.

### Mouse tissue distribution

In previous studies we have shown that there is variation in the expression of the 9-*O*-, 7,9-*O*- and 4-*O*-Ac Sia (15). To better understand the type of variation seen in other tissues, we examined the distribution of those forms, as well as Neu5Gc, in the different tissues of wildtype C57BL/6 mice, as well as those that are CMAH^-/-^. Using the HE-Fc virolectin probes, *O-*acetyl modified Sia were found only in specific tissues and on certain cell subpopulations. Higher staining was seen for cells and mucus in the respiratory and GI tracts, with trachea and colon having particularly strong staining for *O-*acetyl Sia on epithelial cells and associated mucus layers. The percentages of *O-*acetyl modified Sia in the tissues of mice varied widely, with high levels of 7,9-*O*-Ac_2_ and 9*-O-*Ac being found on erythrocytes (∼47%) and the colon (∼19%). The levels in most other tissues varied between 2 and 9%, although these would likely be associated with higher levels on the smaller subsets of cells that were positive for staining with the probes. Neu5Gc was also present in many wild-type mouse tissues with expression varying widely (between 10 and 80%). This is consistent with staining for Neu5Gc previously reported in some mouse tissues (44). 4-*O-*Ac Sia was found in small quantities in only a few tissues, with the highest being found in small intestine (∼2%). This is consistent with previous findings of 4-*O-*Ac levels in mouse brain and liver quantified by GC/MS, although gut associated levels of 4-*O-*Ac were much higher in that study, possibly due to differences in tissue preparation (67). Considering the high level of apparent 4-*O-*Ac specific staining for the mucus in the colon, further investigation of this modification in mouse colonic mucin would be warranted.

### Variation in saliva and other mucins

To define the variation of modified Sia present in secreted mucus across different animals, we examined saliva as it contains many of the same heavily glycosylated proteins and mucins, including MUC5B, that are present in respiratory mucus (49–51). We found a great deal of variability in both the amounts and types of modified Sia present. Some animals (horses, mice, and cows), had larger amounts of *O-*acetyl Sia in their saliva, while human and dogs had primarily unmodified Neu5Ac forms. Pigs are a natural IAV host, and their saliva contained primarily Neu5Gc, so that pig saliva might inhibit IAV infection. Horses had around 10% 4-*O-*Ac Sia present in their saliva. The 4-*O-*Ac modification has been proposed to be a potent inhibitor of many types of viral and bacterial neuraminidases, and to be the γ-inhibitor of horse serum, where it may be present at high levels on the α-2 macroglobulin protein (56, 65, 68). However, as the gene for the 4-*O*-sialyl acetyltransferase has not yet been identified, little is known about its synthesis, expression, or regulation (22, 23).

It has previously been reported that human colonic mucus is enriched for 7,9*-O-* and 9- *O-*Ac (11, 47). However, human saliva and secreted mucus from respiratory cells contained mostly unmodified Neu5Ac, suggesting different expression of modified Sia in the mucus of the respiratory and gastrointestinal tissues. This may be due to the particular functional characteristics of the microbiome in the GI tract compared to other mucosal sites, where 7,9*-O-* and 9-*O-*Ac on colonic mucus may decrease bacterial degradation of sialylated glycans, perhaps improving mucus integrity (69, 70). Microbiome interactions may also explain the increased *O-*acetyl modifications found on both cow and mouse saliva. Cows are ruminants and as part of their digestion, regurgitate partially digested food from their microbe-rich rumen into their mouth to continue chewing as a cud to extract more nutrients. Somewhat similarly, mice are coprophagic and consume feces, along with associated microbes, to re-digest again to improve absorption of nutrients. This would give both cows and mice the potential for more complex mucus and microbe interactions in both the oral cavity and the gut, thus having *O-*acetyl modifications on sialylated glycans on saliva mucus proteins could be advantageous by preventing degradation. However, further research is needed on the roles of these modifications for the interactions between oral and colonic mucus with the microbiome.

### Effects of modified Sia on influenza viruses

IAV use Sia as their primary receptor for host infection, and the specific linkages of Sia to the underlying glycan chain have long been known to influence host tropism (36). We examined the effects of *O-*acetyl and Neu5Gc modifications on IAV HA binding, NA cleavage, and on infection, and found differences among the IAV strains examined. All the different IAV NA tested showed preferential removal of Neu5Ac over any modified Sia form. Cleavage by N1, N7, and N9 were strongly inhibited by mono-acetylated Sia, while N2 and N3 were less affected. All NA had much lower activity against Neu5Gc, di-acetylated Sia, and particularly against Neu5Gc forms with additional *O-*acetylations. This confirms previous reports showing or suggesting inhibition of NA cleavage in H1N1 and H3N2 strains (38, 42), but revealing that there is wide variation of effects on different NA types. Acetylation and Neu5Gc modifications were also inhibitory for HA binding, with soluble HA-Fc sourced from different H1N1 and H3N2 IAV strains showing significantly lower binding to these forms compared to unmodified Neu5Ac in an ELLA assay.

As IAV infects cells at mucosal surfaces in both avian and mammalian hosts, it is likely that viruses interact directly with mucus both to initiate infection and during viral release. Using untreated or esterase treated BSM (to remove 7,9-*O*- and 9-*O*-Ac) it was seen that the unmodified Neu5Ac was most inhibitory, suggesting that *O-*acetyl Sia reduces HA binding to the mucin, allowing binding to the unmodified Sia on the cell surface. Esterase treatment, therefore, allows more efficient binding to the BSM and greater inhibition of IAV infection. For some viruses there is likely a complementarity between the inhibitory effects of 7,9-*O-* and 9-*O-*Ac on HA and NA, where lower HA binding allows the viruses to avoid binding to Sia forms that NA cannot remove efficiently. This effect has been reported for virus grown in the presence of other NA inhibitors (59, 71), and is seen in the balance between the activities of the HA and NA for the α2–3 and α2–6-linked Sia (71, 72). Incubation of virus with the wild-type mouse serum (>80% Neu5Gc) compared to the serum of CMAH^-/-^ mice (100% Neu5Ac) also showed varying effects. Inhibition by WT serum was seen only for pH1N1 virus, while Victoria H3N2 and PR8 H1N1 were inhibited by both sera. This could confirm previous findings that the density of Sia on these serum proteins is the strongest inhibitory factor rather than the type of modified Sia present, as shown for incubation with horse serum (73). However, the variable results for different viruses may indicate variation in sensitivity to Neu5Gc between strains that requires further investigation. It would also be worthwhile comparing inhibition of IAV strains adapted to different host species, particularly species with higher levels of modified Sia present in their respiratory tracts such as horses and pigs.

In summary we have shown that both the *O-*acetyl and Neu5Gc modifications present on secreted glycoproteins in mucus and saliva, as well as on erythrocytes, vary greatly between different species. Some of these modifications inhibit HA binding and NA cleavage, but with a significant variability between IAV strains. While the presence of these modifications can inhibit infection, how they affect virus host tropism and evolution is likely complex and still not fully understood.

## MATERIALS AND METHODS

### Cells and virus

Canine MDCK-NBL2 (ATCC) and A549 (ATCC) cells were grown in Dulbecco’s Modified Eagle Medium (DMEM) with 5% fetal bovine serum and 50 µg/ml gentamicin. Influenza A virus strains pH1N1 (A/California/04/2009), Victoria H3N2 (A/Victoria/361/2011), and PR8 (A/Puerto Rico/8/1934) were rescued from reverse genetics plasmids using previously established protocols (74). Rescued virus were grown to low passage on MDCK-NBL2 cells using infection media containing DMEM, 0.03% BSA, and 1ug/ml TPCK-treated trypsin.

### Erythrocytes, mucus, and saliva

Chicken, cow, sheep, guinea pig, and pig erythrocytes were sourced from Lampire Biological Laboratories (Pipersville, PA). Horse and mouse blood were sourced from The Baker Institute for Animal Health (Cornell University, Ithaca, NY). Dog blood was sourced from the Cornell Veterinary Hospital Diagnostic Center (Ithaca, NY). All erythrocytes were washed in PBS three times and diluted to 5% v/v in PBS. Bovine submaxilliary mucin was purchased from Sigma-Aldrich (St. Louis, MO). Animal studies were all subject to approved protocols from the Cornell Institutional Animal Care and Use Committee.

Saliva from humans was collected by passive drooling following the protocol approved by the University at Buffalo Human Subjects IRB board (study # 030–505616). Informed consent was obtained from all human participants. Saliva from animals was collected by suction using commercially available devices containing absorbent sponges in a syringe-like receptacle (Super-SAL and Micro-SAL, Oasis Diagnostics, Vancouver, WA). Saliva from mice (laboratory strain C57BL/10SnJ) was kindly provided by Jill Kramer (University at Buffalo) using a collection procedure as previously described (75). Saliva from dogs, cows, horses, and pigs, was provided by Erin Daugherity and Luce E. Guanzini (Cornell University). Animals were not allowed to eat or drink prior to the collection to ensure the oral cavity was free of food and other debris. The collection was performed using a commercially available device (Micro-SAL). Large animals were gently restrained and a larger collection device (Super-SAL) was placed under the tongue for up to three minutes, or until fully soaked. Saliva from castrated domestic pigs were also provided by Anja Globig (Friedrich-Loeffler-Institut, Insel Riems - Greifswald, Germany). Saliva from dogs was also kindly provided by Barbara McCabe (Buffalo, NY).

Normal human bronchial epithelial cells (Lonza; cat#CC-2540S) were seeded onto human placental collagen-coated permeable transwell inserts at a density of 2.5×10^4^ cells per well and cultured in bronchial epithelial cell growth basal medium (BEBM) supplemented with bronchial epithelial cell growth SingleQuots (BEGM) in both the apical and basal compartments. After reaching confluence at approximately 7 days post seeding, apical media was aspirated, and basal media replaced with air-liquid interface media containing half DMEM, half BEBM, plus the BEGM SingleQuots to complete differentiation. Apical surfaces of NHBE cells were washed twice with phosphate buffered saline (PBS) to collect mucins for HPLC analysis.

Conditioned media from A549 cells was prepared by washing a fully confluent flask of cells to remove any serum and allowing the cells to grow in serum-free media for 5-7 days. Conditioned media was collected, dialyzed with three volumes of PBS, and concentrated using a 30 kD centrifugal filter (Pall Corporation). Protein concentration was determined using a Qubit 4 fluorometer (Invitrogen).

### Immunohistochemistry of mouse tissues

Expression of O-acetyl modified Sia in various tissues of mice was examined by preparing frozen sections of optimal cutting temperature compound (OCT)-embedded tissue. After a 30 min fixation in 10% buffered formalin, sections were incubated with recombinant virolectins made by expressing nidovirus HE glycoprotein fused to the Fc region of human IgG1, as described by others (16). Nidovirus HEs are specific for *O*-acetyl Sia modifications: MHV-S for 4-*O*-Ac, BCoV-Mebus for 7,9-*O*-Ac (and low recognition of 9-*O*-Ac), PToV-P4 for 9-*O*-Ac. Virolectins were then detected using a biotin-conjugated αFc region secondary antibody followed by incubation with the Vectastain ABC reagent and NovaRed substrate (Vector). Sections were counterstained with hematoxylin.

### Quantification of sialic acid by HPLC

The sialic acid composition from tissues, mucin, and erythrocytes were analyzed by HPLC analysis as previously described (45, 76). In brief, Sia from 20-30 mg of tissue, 50 µg of mucin or saliva, or 100 µl of washed 5% v/v erythrocytes were release using 2M acetic acid at 80°C for 3 hr followed by filtration through a 10kD centrifugal filter (Microcon) and dried using a vacuum concentrator (SpeedVac). Released Sia were labeled with 1,2-diamino-4, 5-methylenedioxybenzene (DMB, Sigma Aldrich) for 2.5 hr at 50°C. HPLC analysis was performed using a Dionex UltiMate 3000 system with an Acclaim C18 column (ThermoFisher) under isocratic elution in 7% methanol, 7% acetonitrile, and 86% water. Sia standards were bovine submaxillary mucin, normal horse serum, and commercial standards for Neu5Ac and Neu5Gc (Sigma Aldrich). Pre-treatment of samples with 30 µg/ml esterase-active BCoV HE-Fc overnight at 37°C removed 7,9-*O*- and 9-*O-*acetyl modifications. Final data analysis was completed using PRISM software (GraphPad, version 8).

### Biotinylated a2–6-linked sialosides

Biotinylated α2–6-linked sialosides Sia α2– 6LacNAc-biotin containing Neu5Ac, Neu5Gc, or Neu5Ac9NAc as the sialic acid form were synthesized from LacNAc-biotin (77) as the acceptor substrate and Neu5Ac, ManNGc (78), or ManNAc6NAc (62) as the donor precursor using a one-pot multienzyme sialylation system similar to that described previously (79).

### IAV HA affinity for 9-*O*-Ac modified Sia

HA-Fc constructs were produced as previously described (80). HA-Fc binding to sialosides was performed as previously described (63, 64). In brief, ELISA-grade 96 well plates (ThermoFisher Scientific) were coated with 5 µg of HA-Fc for overnight at 4°C. Plates were then washed 3× with PBS and blocked using 1x Carbo Free Blocking Buffer (Vector Labs, Burlingame, CA) for 1 hr. After blocking, plates were washed once with PBS and treated with sialosides diluted in PBS for 3 hr at room temperature, then washed 3× with PBS. Plates were then incubated with HRP-streptavidin complex (Vector Labs) for 45 min, washed 3× with PBS, then incubated with 3,3’,5,5’-tetramethyl benzidine (TMB, Thermo Fisher Scientific). TMB development was stopped with 2M sulfuric acid and then analyzed using a colorimetric plate reader (Multiskan EX, ThermoFisher Scientific). Data analysis was completed in PRISM software (GraphPad, version 8).

### Generation of NA VLPs

NA sequences were obtained from GenBank (N1: ACP44181, N2: AGC70842, N3: ACL11962, N7: AAR11367, N9: ARG43209). Sequences were codon optimized, tagged, and ordered through Biomatik in the pcDNA3.1(+) vector. To produce VLPs, HEK-293T cells were seeded in 15cm plates and transfected when 80% confluent. Cells were transfected with 4 µl of Polyethylenimine (PEI) (Polysciences cat# 23966-2) at 1 mg/ml concentration for every 1 µg plasmid DNA stock in 9ml of Opti-MEM. Eight hours post transfection, 6ml of pre-warmed Opti-MEM was added. Supernatant was collected 72 hours post transfection and purified using ultracentrifugation (110,000xg, 1.5 hours, 4°C) through a 20% sucrose cushion, then the pellet re-suspended in PBS and stored at 4°C.

### NA cleavage assay with NA VLPs

Bovine sub-maxillary mucin (BSM) or erythrocytes from horses and chickens were used as a substrate to determine NA activity of the different NA VLPs. Briefly, 50 µg of BSM or 5% v/v washed erythrocytes in PBS were treated with 1:100 NA VLPs or 1:100 *Arthrobacter ureafaciens* NA (NeuA, New England BioLabs) for 4 hours at 37°C. Free Sia was collected and prepared for HPLC analysis as above.

### Mucin and serum inhibition of infection

MDCK cells were seeded to ∼80% confluency in 12 well plates with cells allowed to settle for 6 hrs. Virus was diluted in PBS to give MOI of 0.5 and then mixed with 20 µg untreated BSM or esterase treated BSM and incubated for 45 min at room temp. For mouse serum inhibition, virus was mixed with 200 µg wild-type or CMAH^-/-^ serum instead. Serum- or mucin-treated virus was then added to washed MDCK cells and incubated for 1 hr with tilting to prevent cell drying. Inoculum was then removed, media added and cells were incubated for 10 hrs. Cells were then harvested, stained with an anti-influenza A NP antibody, and analyzed for infection using a Millipore Guava EasyCyte Plus flow cyotometer (EMD Millipore, Billerica, MA) with analysis using FlowJo software (TreeStar, Ashland, OR). Statistical analyses were performed in PRISM software (GraphPad, version 8).

## ACKNOWLEDGEMENTS

We thank Wendy Weichert for expert technical support. We also thank Lubov Neznanova and Jill M. Kramer (University of Buffalo), along with Anja Globig (Friedrich Loeffler Institute, Germany), for sharing samples of saliva.

---

**Supplemental Figure 1.**
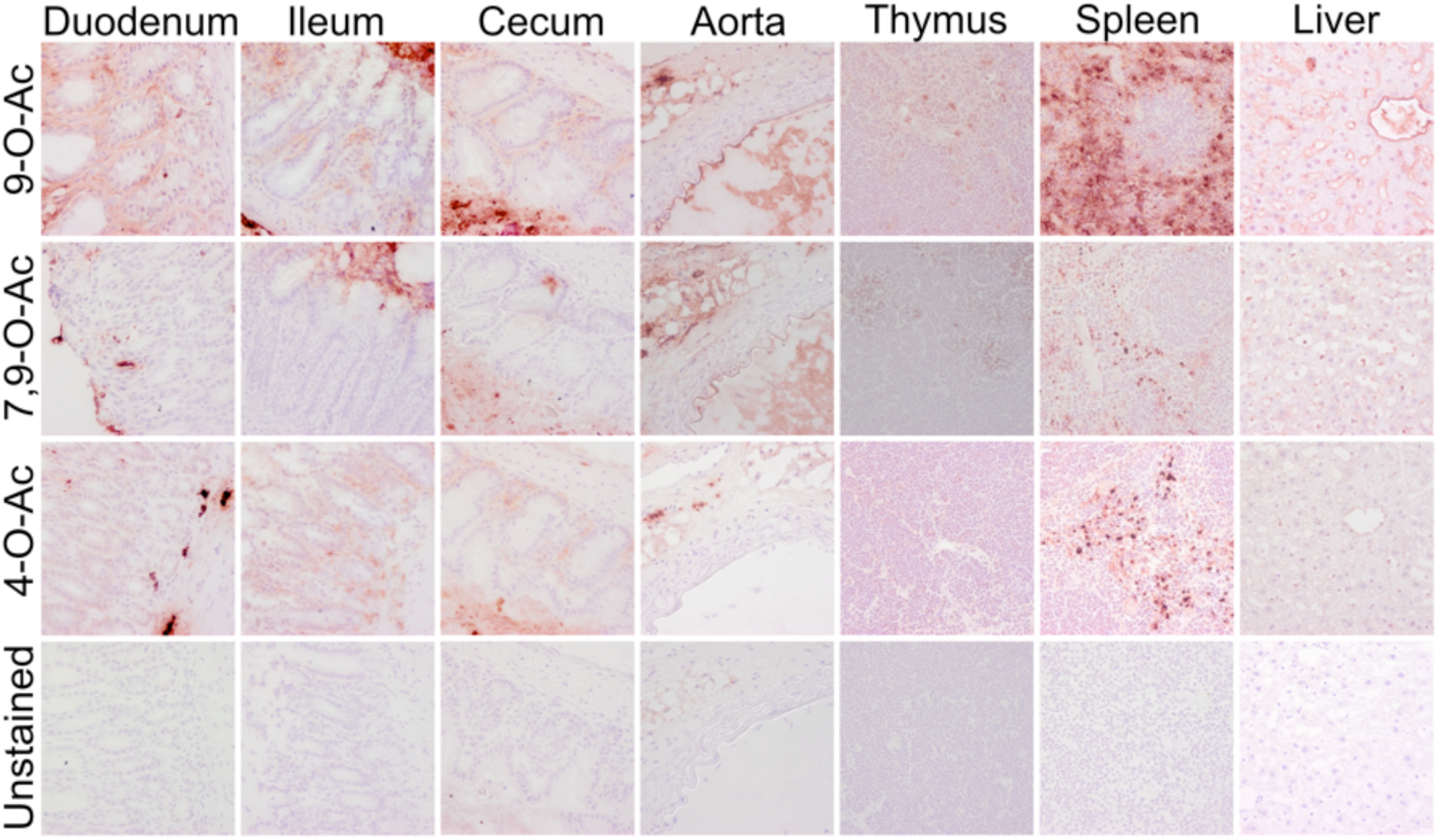
Expression of *O*-acetylated Sia varies between tissues in wild-type C57BL/6 mice Frozen tissue sections were stained using virolectins derived from the hemagglutinin esterases (HE-Fc) of various nidoviruses with high specificity for the different *O-*acetyl modified Sia forms. Sections were counterstained with hematoxylin and imaged at 40x magnification

**Supplemental Table 1.** Average relative Sia quantities as determined by HPLC analysis for each tissue tested in WT C57BL/6 mice (n=3). Table shows the proportion of total Sia for each variant given as a percentage, with the sum of all *O-*acetyl forms combined given in the far right column. If a Sia form was below detection, this is indicated by b/d.

| Tissue | Neu5Gc | Neu5Ac | Neu5,9Ac | Neu5,7Ac2 | Neu5Gc8Ac | Neu4,5Ac2 | Neu5,8Ac2 | Neu5,7,8/9Ac3 | O-Ac sum |
| --- | --- | --- | --- | --- | --- | --- | --- | --- | --- |
| Lung | 50.2 | 47.6 | 1.2 | b/d | 1.3 | 0.5 | 0.2 | 0.4 | 3.6 |
| Trachea | 43.0 | 53.5 | 1.3 | b/d | 1.5 | 0.4 | 0.2 | 0.4 | 3.7 |
| Stomach | 27.2 | 68.8 | 2.0 | 0.7 | 0.6 | 0.7 | 0.2 | b/d | 4.3 |
| Aorta | 58.4 | 38.2 | 1.4 | b/d | 1.7 | b/d | 0.2 | b/d | 3.3 |
| Brain | 9.8 | 83.1 | 4.5 | 1.0 | 2.5 | 1.1 | 0.5 | b/d | 9.5 |
| Esophagus | 53.1 | 43.2 | 1.5 | b/d | 1.4 | 0.6 | 0.2 | 0.2 | 3.9 |
| Spleen | 49.6 | 45.2 | 2.4 | b/d | 1.0 | 1.1 | 0.3 | 0.8 | 5.6 |
| Testes | 57.1 | 40.4 | 1.1 | b/d | 0.8 | 1.2 | b/d | b/d | 3.0 |
| Salivary Gland | 11.1 | 85.5 | 1.9 | 3.6 | 0.2 | 0.2 | 0.1 | b/d | 6.0 |
| Liver | 72.6 | 23.8 | 1.0 | b/d | 1.7 | 0.8 | 0.2 | b/d | 3.7 |
| Small Intestine (duodenum) | 27.6 | 68.7 | 1.3 | b/d | 0.6 | 1.8 | 0.1 | b/d | 3.8 |
| Uterine Horn | 45.3 | 52.4 | 0.8 | b/d | 1.4 | 0.4 | 0.1 | b/d | 2.6 |
| Kidney | 43.8 | 53.5 | 1.1 | 0.5 | 1.0 | 0.4 | 0.1 | 0.2 | 3.4 |
| Tongue | 51.5 | 45.8 | 1.2 | b/d | 1.1 | 0.5 | n/d | b/d | 2.8 |
| Heart | 43.2 | 53.3 | 1.8 | b/d | 0.9 | 0.8 | 0.2 | 0.6 | 4.3 |
| Large Intestine (colon) | 5.9 | 77.2 | 9.0 | 3.8 | b/d | b/d | 1.6 | 2.6 | 16.9 |
| Thymus | 52.1 | 44.8 | 1.1 | b/d | 0.6 | 1.3 | 0.1 | b/d | 3.1 |
| Pancreas | 38.5 | 59.1 | 1.2 | b/d | 0.7 | 0.4 | 0.2 | b/d | 2.5 |

**Supplemental Table 2.**
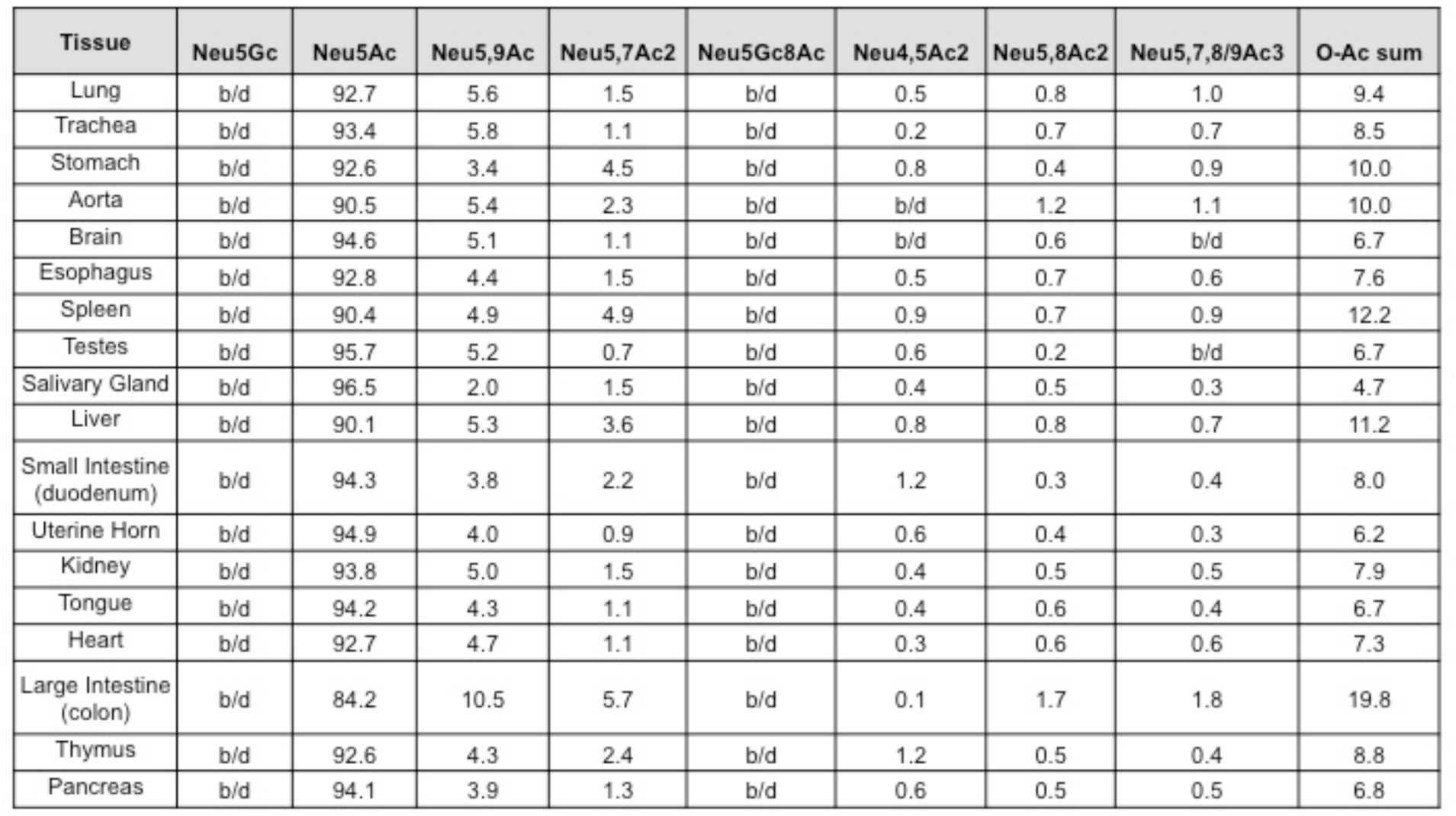
Average relative Sia quantities as determined by HPLC analysis for each tissue tested in CMAH^-/-^ C57BL/6 mice (n=3). Table shows the proportion of total Sia for each variant given as a percentage, with the sum of all *O-*acetyl forms combined given in the far right column. If a Sia form was below detection, this is indicated by b/d.

| Tissue | Neu5Gc | Neu5Ac | Neu5,9Ac | Neu5,7Ac2 | Neu5Gc8Ac | Neu4,5Ac2 | Neu5,8Ac2 | Neu5,7,8/9Ac3 | O-Ac sum |
| --- | --- | --- | --- | --- | --- | --- | --- | --- | --- |
| Lung | b/d | 92.7 | 5.6 | 1.5 | b/d | 0.5 | 0.8 | 1.0 | 9.4 |
| Trachea | b/d | 93.4 | 5.8 | 1.1 | b/d | 0.2 | 0.7 | 0.7 | 8.5 |
| Stomach | b/d | 92.6 | 3.4 | 4.5 | b/d | 0.8 | 0.4 | 0.9 | 10.0 |
| Aorta | b/d | 90.5 | 5.4 | 2.3 | b/d | b/d | 1.2 | 1.1 | 10.0 |
| Brain | b/d | 94.6 | 5.1 | 1.1 | b/d | b/d | 0.6 | b/d | 6.7 |
| Esophagus | b/d | 92.8 | 4.4 | 1.5 | b/d | 0.5 | 0.7 | 0.6 | 7.6 |
| Spleen | b/d | 90.4 | 4.9 | 4.9 | b/d | 0.9 | 0.7 | 0.9 | 12.2 |
| Testes | b/d | 95.7 | 5.2 | 0.7 | b/d | 0.6 | 0.2 | b/d | 6.7 |
| Salivary Gland | b/d | 96.5 | 2.0 | 1.5 | b/d | 0.4 | 0.5 | 0.3 | 4.7 |
| Liver | b/d | 90.1 | 5.3 | 3.6 | b/d | 0.8 | 0.8 | 0.7 | 11.2 |
| Small Intestine (duodenum) | b/d | 94.3 | 3.8 | 2.2 | b/d | 1.2 | 0.3 | 0.4 | 8.0 |
| Uterine Horn | b/d | 94.9 | 4.0 | 0.9 | b/d | 0.6 | 0.4 | 0.3 | 6.2 |
| Kidney | b/d | 93.8 | 5.0 | 1.5 | b/d | 0.4 | 0.5 | 0.5 | 7.9 |
| Tongue | b/d | 94.2 | 4.3 | 1.1 | b/d | 0.4 | 0.6 | 0.4 | 6.7 |
| Heart | b/d | 92.7 | 4.7 | 1.1 | b/d | 0.3 | 0.6 | 0.6 | 7.3 |
| Large Intestine (colon) | b/d | 84.2 | 10.5 | 5.7 | b/d | 0.1 | 1.7 | 1.8 | 19.8 |
| Thymus | b/d | 92.6 | 4.3 | 2.4 | b/d | 1.2 | 0.5 | 0.4 | 8.8 |
| Pancreas | b/d | 94.1 | 3.9 | 1.3 | b/d | 0.6 | 0.5 | 0.5 | 6.8 |

